# Assessment of mapping strategies for determining the 5□-end of mRNAs and long-noncoding RNAs with short read sequences

**DOI:** 10.1101/2020.03.14.982991

**Authors:** Shuhei Noguchi, Hideya Kawaji, Takeya Kasukawa

**Affiliations:** RIKEN Center for Integrative Medical Sciences, Yokohama, Kanagawa 230-0045, Japan; RIKEN Preventive Medicine and Diagnosis Innovation Program, Wako, Saitama 351-0198, Japan; Institute for Protein Research, Osaka University, Suita, Osaka 565-0871, Japan

**Keywords:** Transcription Start Site, CAGE, Mapping, Transcriptome, Reference transcriptome

## Abstract

**Background:** Genome mapping is an essential step in data processing for transcriptome analysis, and many previous studies have evaluated various methods and strategies for mapping RNA-seq data. Cap Analysis of Gene Expression (CAGE) is a sequencing-based protocol particularly designed to capture the 5□-ends of transcripts for quantitatively measuring the expression levels of transcription start sites genome-wide. Because CAGE analysis can also predict the activities of promoters and enhancers, this protocol has been an essential tool in studies of transcriptional regulation. Typically, the same mapping software is used to align both RNA-seq data and CAGE reads to a reference genome, but which mapping software and options are most appropriate for mapping the 5□-end sequence reads obtained through CAGE has not previously been evaluated systematically.

**Results:** Here we assessed various strategies for aligning CAGE reads, particularly ∼50-bp sequences, with the human genome by using the HISAT2, LAST, and STAR programs both with and without a reference transcriptome. One of the major inconsistencies among the tested strategies involves alignments to pseudogenes and parent genes: some of the strategies prioritized alignments with pseudogenes even when the read could be aligned with coding genes with fewer mismatches. Another inconsistency concerned the detection of exon-exon junctions. These preferences depended on the program applied and whether a reference transcriptome was included. Overall, the choice of strategy yielded different mapping results for approximately 2% of all promoters.

**Conclusions:** Although the various alignment strategies produced very similar results overall, we noted several important and measurable differences. In particular, using the reference transcriptome in STAR yielded alignments with the fewest mismatches. In addition, the inconsistencies among the strategies were especially noticeable regarding alignments to pseudogenes and novel splice junctions. Our results indicate that the choice of alignment strategy is important because it might affect the biological interpretation of the data.

## Background

Next-generation sequencing (NGS) has been widely used for transcriptome analysis, for example RNA sequencing (RNA-seq). Many experimental protocols for transcriptome analysis have been developed and applied. An essential step in the data processing for transcriptome analysis is genome mapping, in which sequence reads are aligned to a reference genome and the alignment results are used to quantify gene-level or transcript-level sequence expression. Several publicly available programs have been developed for genome mapping of RNA-seq data. These programs use various algorithms to align sequences with splice junctions and offer various options, for example, whether they use a reference transcriptome like GENCODE [1] and RefSeq [2] to predict the splice junctions. Because mapping typically consumes the majority the time in required for data processing and mapping results can influence subsequent data analysis, many studies have compared various mapping programs and options to evaluate which combination is most appropriate for processing RNA-seq data [3–5].

For one variation of transcriptome analysis by NGS, several protocols—such as, Cap Analysis of Gene Expression (CAGE) [6, 7], Transcription Start Site sequencing (TSS-seq) [8, 9], and RNA Annotation and Mapping of Promoters for Analysis of Gene Expression (RAMPAGE) [10]—selectively capture the 5□-end of transcripts for comprehensive analysis of transcription start sites (TSSs) at 1-bp resolution. These protocols accurately identify the locations of the 5□-end of expressed transcripts and their genome-wide expression levels, enabling us to estimate the transcriptional activity of promoters [11–13] and even enhancers [14]. In this regard, the Functional ANnoTation of the Mammalian Genome 5 (FANTOM5) project [13] involved comprehensive CAGE analysis of more than 1800 human samples of primary cells, cell lines, tissues, and time-course transitions during cell differentiation. This effort revealed the presence of approximately 200,000 TSSs (that is, promoters) and approximately 65,000 enhancers in the human genome, and produced a comprehensive atlas of human promoters and enhancers [15, 16].

Given that each CAGE read is a part of an RNA sequence, it might seem that the same genome mapping protocols used for RNA-seq could potentially be used for mapping CAGE reads. However, in actuality, mapping requirements differ between RNA-seq and CAGE data. One difference regards the use of reference transcriptomes. In previous studies [13, 17, 18], CAGE reads were mapped without reference transcriptomes, for the following reasons. Public reference transcriptomes were (and are) not always complete at the 5□-end of transcribed regions [13]—a situation that affects the genomic mapping of CAGE reads. Moreover, the CAGE reads were relatively short—shorter than the first exon—and therefore lacked splice junctions.

The mapping protocols for RNA-seq and CAGE data also differ because the FANTOM5 project used an original mapping software (delve) for CAGE data due to the extremely short read lengths and high error rates in the HeliScope sequencer [13]. As a consequence of addressing those issues, this software cannot support alignments containing splice junctions.

Another key difference regarding mapping requirements for RNA-seq compared with CAGE data is that the purpose of CAGE analysis is to precisely identify the 5□-end of transcripts. This goal necessitates complete alignment of the 5□-ends of CAGE sequence reads, which is not always necessary for RNA-seq.

Owing to these different requirements, the necessary features and components of software protocols for CAGE read mapping must be evaluated independent from those for RNA-seq data. However, no currently available systematic evaluation focuses particularly on which mapping programs and options to choose for genome mapping of CAGE reads. In addition, minimal sequence read lengths have become longer in recent protocols (e.g., approximately 50 bp in nAnTi-CAGE [19]), but in many transcripts, the first exons are shorter than 50 bp [20]. Consequently, whether the introduction of known splice junctions in the reference transcriptomes actually improves the alignment of CAGE reads must be reassessed. Another consideration is that, in a process known as ‘soft clipping,’ recent genome mapping programs disregard (clip) mismatched 5□- and 3□-end bases in sequence reads rather than force these sequences to map with mismatches. Because the exact positions of the 5□-ends of sequence reads are important in CAGE data analysis, soft clipping has the potential to undermine the results. Therefore, an independent comparison and evaluation of mapping software and options, with specific focus on CAGE reads, is urgently needed.

In this study, to investigate which method is most appropriate for the genome mapping of CAGE reads, we compared the mapping results we obtained by using several high-throughput protocols for RNA-seq—that is, HISAT2 [5], LAST [21], and STAR [4]— both with and without the inclusion of a reference transcriptome. In our evaluation, we especially focused on genes with short first exons, pseudogenes, mapped reads including novel splice junctions, and promoter-level expression values.

## Results

### Selection of data set and alignment software

HISAT2 and STAR are widely used splice-aware short-read aligners for RNA-seq data [4, 5]. We ran these programs under two conditions: (i) with a reference transcriptome model from the GENCODE dataset (termed ‘HISAT2-kss’ and ‘STAR-kss’; the tag kss indicates that known splice sites are incorporated into the mapping strategy) and (ii) without reference transcriptome models (termed ‘HISAT2-nss’ and ‘STAR-nss’; nss: known splice sites are omitted from the analysis). We also evaluated LAST (termed ‘LAST-nss’), a program that can be used for spliced alignments containing many mismatches and gaps and that has been used recently for mapping of long-read sequencing [21].

We used the CAGE data set obtained from the TC-YIC cell line, a cell line derived from human non-islet cell insulinoma cells (DRA002420) [17], as a test set to evaluate of the mapping strategies. Data from 93 samples comprising biological triplicates were obtained by using the nAnT-iCAGE (no-amplification non-tagging CAGE) protocol [19], which is a variation of the CAGE protocols developed for the Illumina sequencer. The read length was 51 bp with a single end, consisting of 3-bp sample indexes and 48-bp cDNA reads corresponding to the 5□-end of RNAs. The CAGE dataset obtained from the TC-YIC cell line is typical of the large-scale human CAGE datasets generated by using the nAnT-iCAGE protocol.

### Summary statistics of all mapping results

We summarized the mapping results from the five strategies to calculate the numbers of mapped reads, promoter rates, numbers of expressed promoters, detected known splice junctions, detected novel splice junctions, and expressed genes (Fig. 1). In the promoter level analysis, we used the FANTOM5 CAGE peaks [16] as a reference set of promoters. We noted that mapped reads with unaligned regions larger than 5 bp at the 5□-ends of alignments were flagged as ‘unmapped’ after running the alignment programs; such reads might not represent true 5□-ends of transcripts, thus potentially affecting quantification of TSS expression.

**Figure 1.**
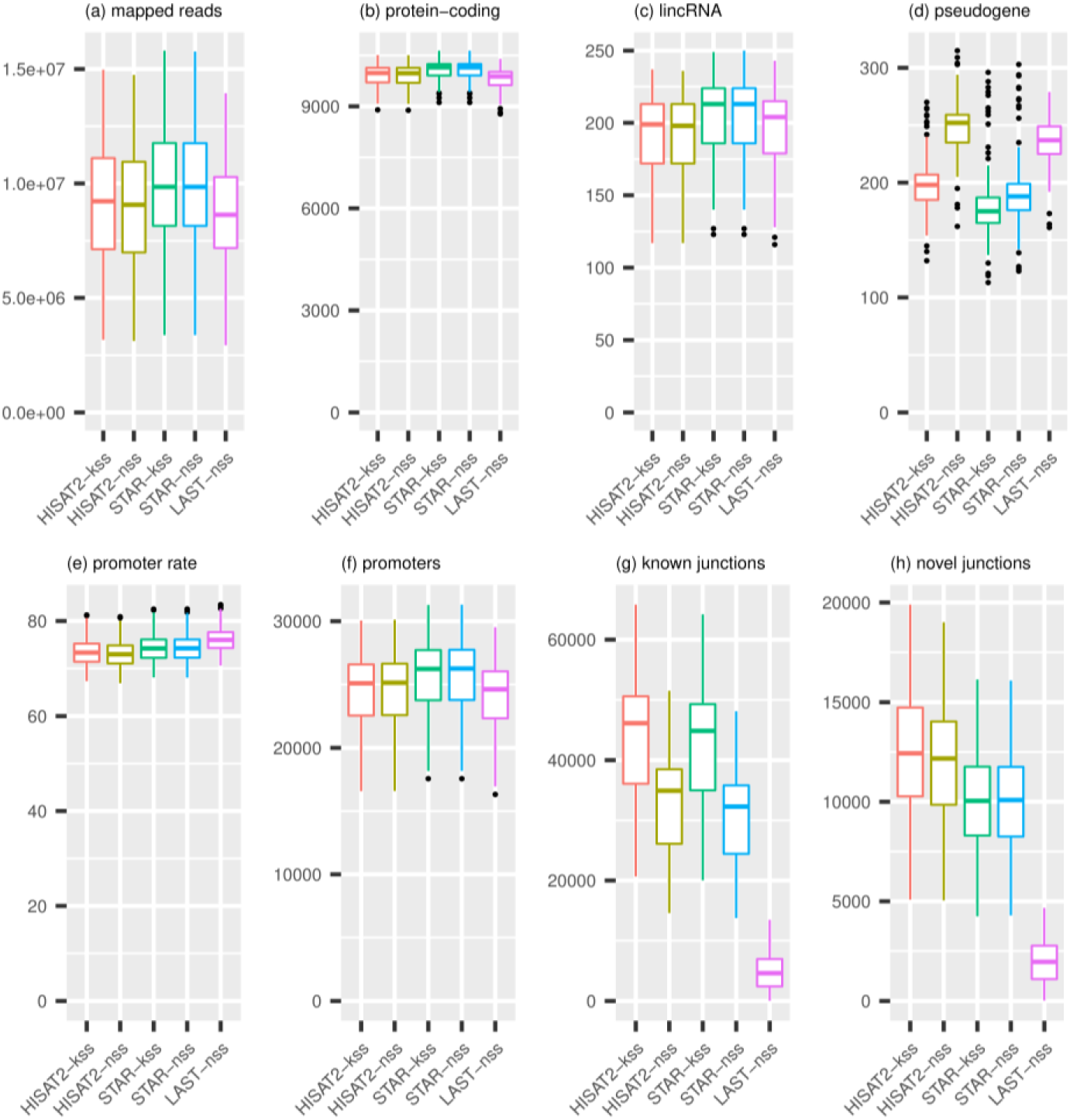
Mapping summary. Boxplots of (a) mapped reads; number of expressed genes associated with (b) protein-coding genes, (c) lincRNAs, and (d) pseudogenes; (e) promoter rate (%); (f) number of promoters; (g) number of detected known splice junctions; and (h) number of detected novel splice junctions are shown for each mapping condition.

The numerical difference between the number of mapped reads with the reference transcriptome and without it (that is, STAR-kss □ STAR-nss and HISAT2-kss □ HISAT2-nss) was less than ones between difference mapping programs (Fig. 1a). In regard to the numbers of mapped reads and expressed promoters, both STAR-kss and STAR-nss yielded higher values than the other three strategies (Fig. 1a, c). Interestingly, LAST-nss was associated with a low mapping rate but a high promoter rate (Fig. 1a, e). This pattern might relate to our observation that CAGE reads that were mapped to non-promoter regions by the other four mapping strategies tended to remain unmapped in LAST-nss (explained in the later section).

Using the FANTOM5 classification of promoters to protein-coding genes, long intervening noncoding RNAs (lincRNAs), and pseudogenes, we calculated the number of expressed genes in each category (Fig. 1b□d). Among all mapping strategies, the numbers of expressed genes associated with protein-coding genes and lincRNAs (Fig. 1b, c) were similar in pattern to those of overall mapped reads (Fig. 1a). However, the number of expressed pseudogenes varied depending on the mapping strategy (Fig. 1d) and followed the order of STAR-kss ≈ STAR-nss < HISAT2-kss < HISAT2-nss < LAST-nss.

### Promoter expression

To evaluate how differences in mapping strategy affect results for promoter-level expression, we calculated the expression levels of FANTOM5 promoters [13, 22] for each mapping strategy and then identified promoters that were significantly differentially expressed among the mapping strategies (Fig. 2). This analysis revealed that at most ∼1.5% of ∼200,000 promoters were identified as differentially expressed between two mapping strategies (Fig. 2a). The correlations between HISAT2-kss and HISAT2-nss and between STAR-kss and STAR-nss were much higher than any method compared with LAST-nss (Fig. 2b); promotor-level expression values tended to be lower for LAST-nss than for the other 4 strategies. For both HISAT2 and STAR, whether a reference transcriptome was included (or not) had little effect on results regarding promotor-level expression (Fig. 2a).

**Figure 2.**
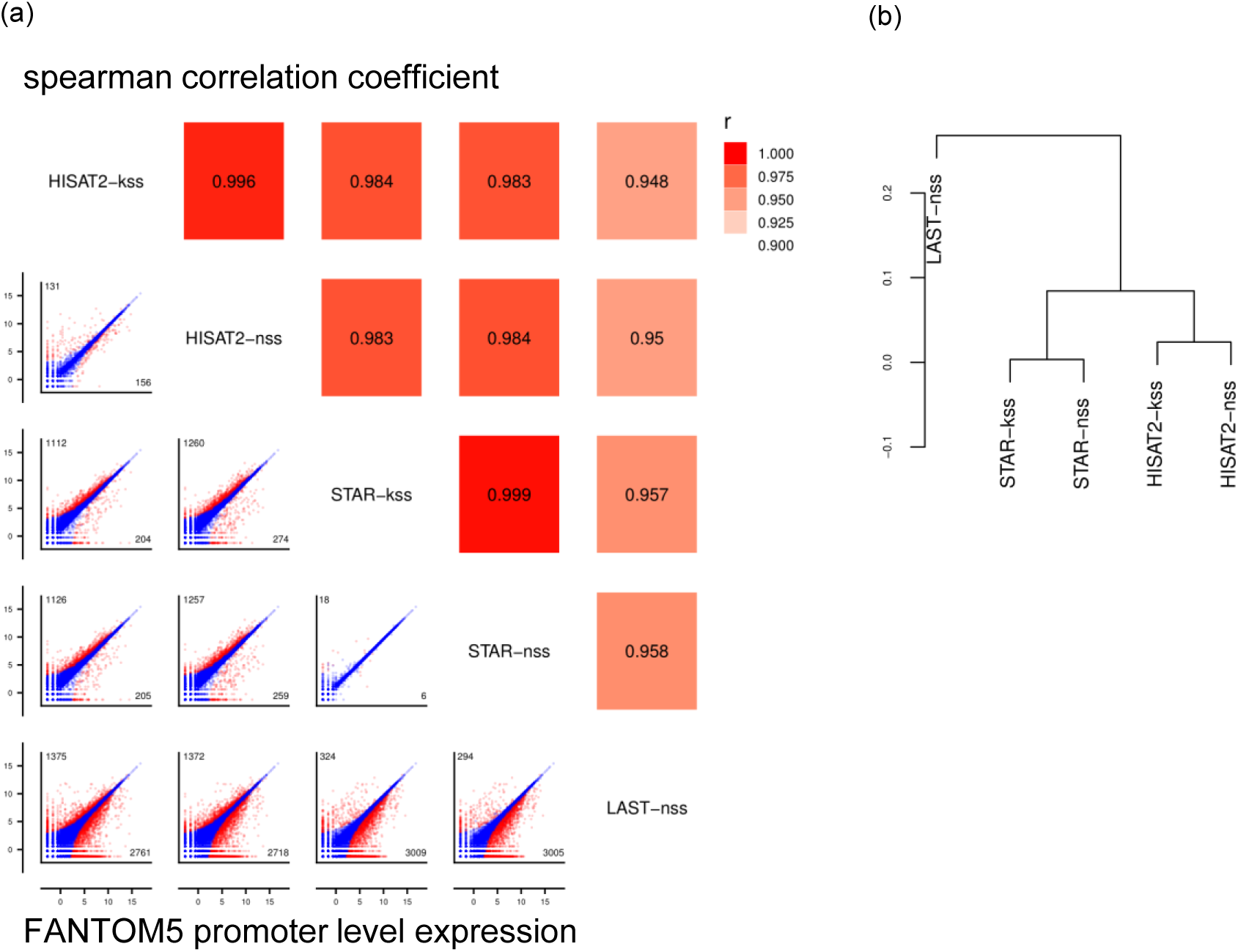
Differences in promoter expression among mapping strategies. a) Scatter plots of promoter expression from pair of mapping strategies using biological triplicate samples (DRR021875, DRR021876, and DRR021877). Dots represent the average promoter expression of the 3 replicates. Red dots indicate significantly differentially expressed promoters (q < 0.01 by edgeR), and blue dots are similarly expressed promoters. The values at the top right are the Spearman correlation coefficients of promoter expression among the triplicate samples. The number at the top left in each scatter plot is the number of significantly highly expressed promoters in the mapping strategy along the *y*-axis compared with that on the *x*-axis. The number at the bottom right in each scatter plot is the number of promoters with low expression. b) Hierarchical clustering of mapping strategies according to promoter expression in the same triplicate samples.

### Inconsistency of mapped reads

To deeply investigate differences among mapping strategies, we next examined reads that aligned to different positions depending on the mapping strategy used (Fig. 3). Mapping inconsistency was lowest between STAR-kss and STAR-nss and maximum between STAR-kss or -nss and LAST-nss (Fig. 3b). The discrepancy primarily involves protein-coding genes and pseudogenes. On average, marked mismatches in inconsistent alignments were fewest in STAR-kss (Fig. 3c), suggesting that more signals of genuine transcription were assigned.

**Figure 3.**
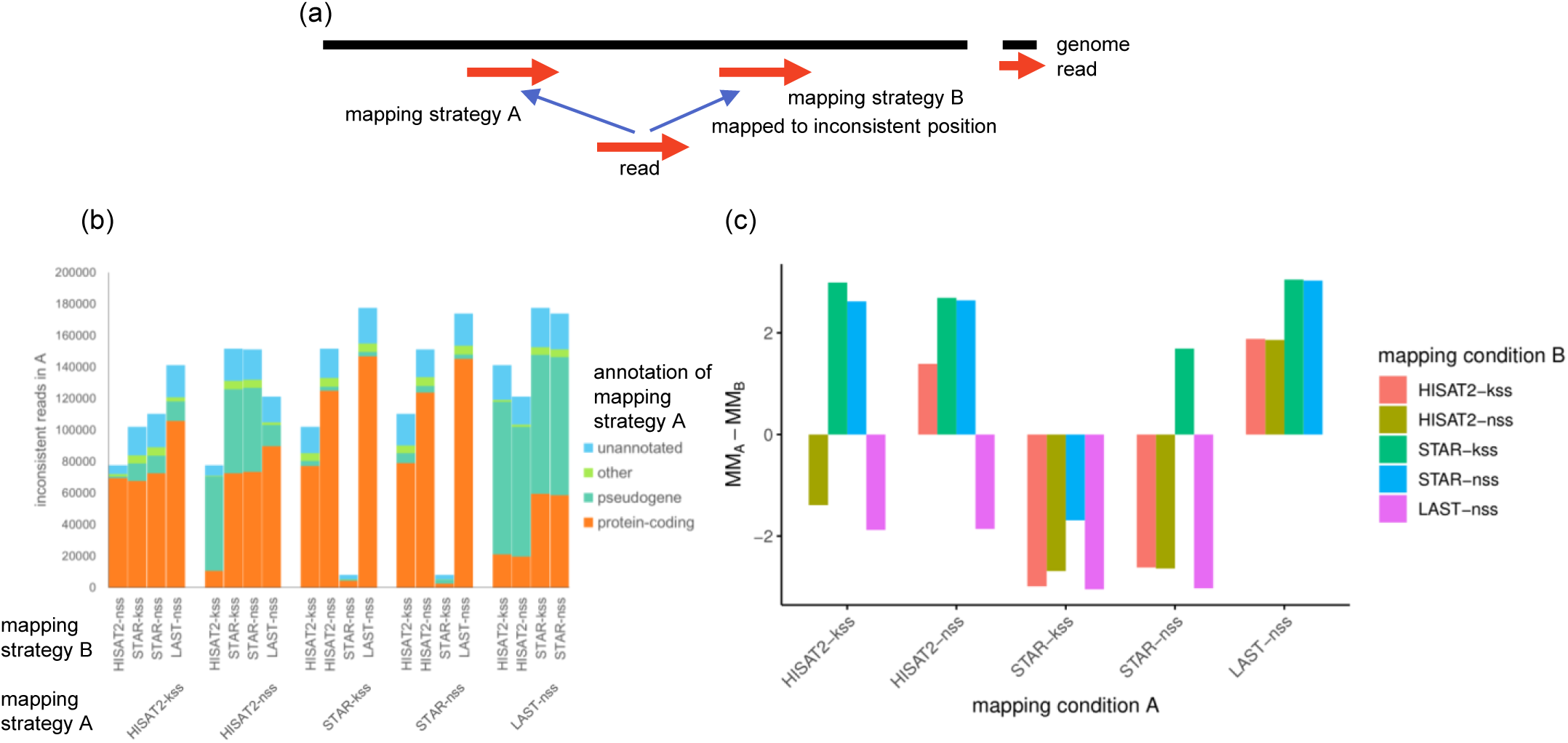
Inconsistently mapped reads with the number of mismatches. a) A model of inconsistently mapped reads. An inconsistently mapped read is one that was mapped to 2 different positions by 2 different mapping strategies (e.g., A and B). b) The number of inconsistently mapped reads between mapping strategies A and B. The reads were classified as protein-coding gene, pseudogene, other type of gene (other), and unannotated according to mapped genes in mapping strategy A. c) The difference in the mean number of mismatches in alignments of inconsistently mapped reads between mapping strategies A and B. The *y*-axis represent Mean(MM_Ai_ – MM_Bi_), where i is an inconsistently mapped read, MM_Ai_ is the number of mismatches for read ‘i’ in mapping strategy A, and MM_Bi_ is the number of mismatches for read ‘i’ in mapping strategy B. A positive value indicates a greater number of mismatches in reads mapped by strategy A compared with strategy B.

### Reads mapped to pseudogenes

As we showed earlier, pseudogenes are one of the main causes of inconsistent alignments between mapping strategies. To confirm this finding, we examined reads that were inconsistently mapped to pseudogenes (Fig. 4 and S1). We first evaluated the number of reads that were mapped to pseudogenes by one mapping strategy but to a different region by the other. Overall, most of reads that were mapped to pseudogenes by HISAT2-kss and STAR-kss were consistently mapped to the same positions by HISAT2-nss and STAR-nss, respectively (Fig. S1). However, about 30% of reads that were mapped to pseudogenes by HISAT2-nss were mapped to different positions by HISAT2-kss (Fig. S1), suggesting that the use of the reference transcriptome greatly facilitated the discovery of parental genes with exon–intron structures. In addition, approximately half of the reads that were aligned to pseudogenes by LAST-nss mapped to different places by the other strategies (Fig. S1). Among the reads that mapped inconsistently between LAST-nss and the other mapping strategies, about half were mapped to pseudogenes in LAST-nss (Fig. 3b), but most of them were mapped to protein-coding genes in the other mapping strategies (Figs. 4c, d). For example, STAR-kss mapped multiple reads to the protein-coding gene RPL13 but LAST-nss mapped the same reads to the RPL13 pseudogene RPL13P12, likely owing to the approach used to accommodate transcriptome reference–free alignments in LAST-nss.

**Figure 4.**
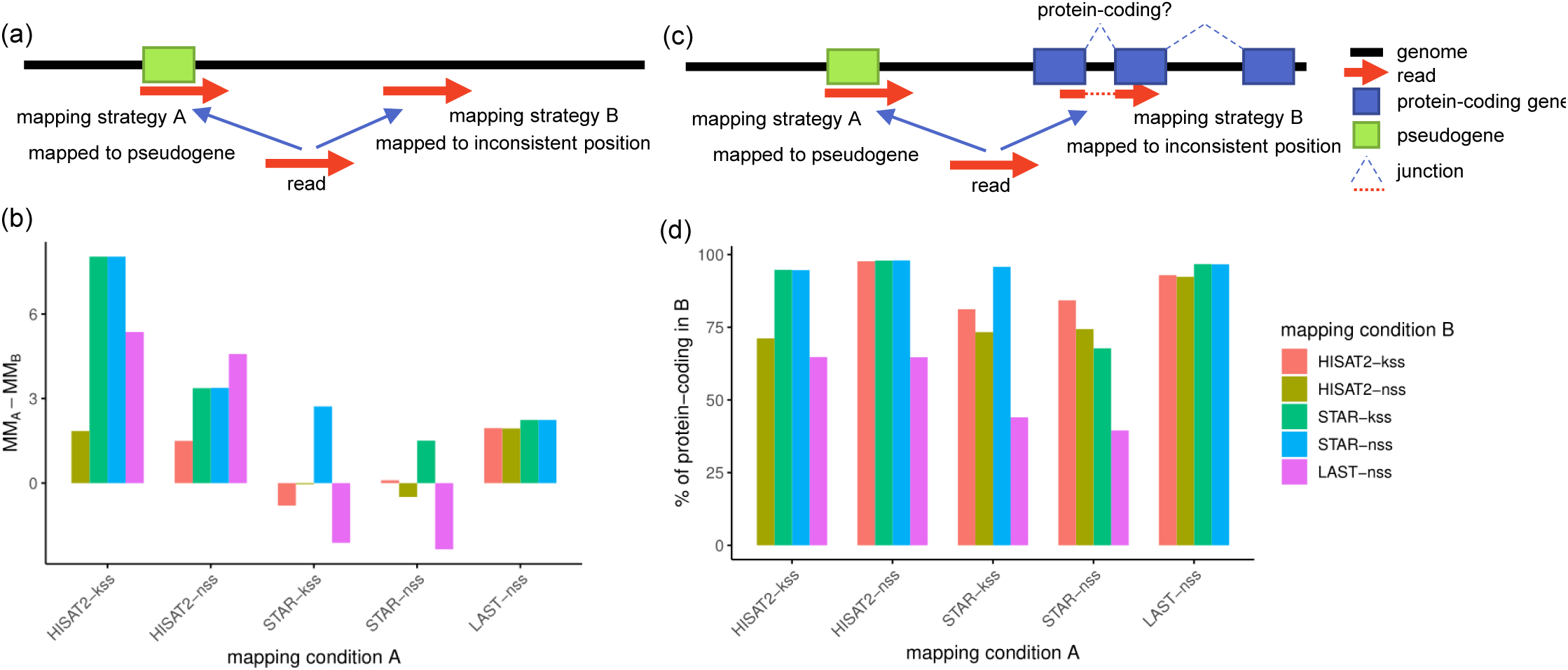
Inconsistently mapped reads to pseudogene. a) A model of an inconsistently mapped read to a pseudogene is one in which a read is mapped to a pseudogene by one strategy (i.e., mapping strategy A) but to an inconsistent position by the other (i.e., mapping strategy B). b) The difference in the mean number of mismatches between mapping strategies A and B. The *y*-axis indicates Mean(MM_Ai_ – MM_Bi_), where ‘i’ is an inconsistently mapped read to a pseudogene, MM_Ai_ is the number of mismatches for read i in mapping strategy A, and MM_Bi_ is the number of mismatches of read i in mapping strategy B. A positive value indicates more mismatches from mapping strategy A than B. c) A model of an inconsistently mapped read to a pseudogene and its parent gene. An inconsistently mapped read to a pseudogene and its parent gene is one in which a read is mapped to a pseudogene by strategy A but to an inconsistent position in a protein-coding gene region by strategy B. d) The percentage of reads that strategy B mapped to protein-coding gene regions relative to all inconsistently mapped reads to pseudogenes and their parent genes.

We next compared the mean number of mismatches in alignments inconsistently mapped to pseudogenes. Alignments mapped to pseudogenes by HISAT2-kss, HISAT2-nss, and LAST-nss had more mismatches than alignments mapped to non-pseudogene regions in the other mapping strategies (Fig. 4b). In contrast, STAR yielded fewer mismatches in alignments mapped to pseudogenes than did the other strategies (Fig. 4b).

### Reads mapped over novel splice junctions

We also analyzed inconsistently mapped reads that included splice junctions, which are another potential cause of mapping inconsistencies (Fig. 5). The number of detected known splice junctions differed substantially depending on whether the reference transcriptome was used (or not), whereas the difference between HISAT2-kss and STAR-kss or between HISAT2-nss and STAR-nss was very small (Fig. 1g). In contrast, the number of novel splice junctions was slightly greater in HISAT2-kss/-nss than STAR-kss/-nss (Fig. 1h) and was lowest in LAST-nss. In addition, the mean number of mismatches in alignments containing novel splice junctions was generally lower than ones without novel splice junctions (Fig. 5b), except for LAST-nss versus STAR-kss and STAR-nss. Furthermore, we found that, except for LAST-nss, most reads inconsistently mapped with novel splice junctions were aligned to different loci without splice junctions (Fig. 5d). However, the reads that were inconsistently mapped with novel splice junctions in LAST-nss were aligned with known splice junctions in another mapping strategy (Fig. 5f).

**Figure 5.**
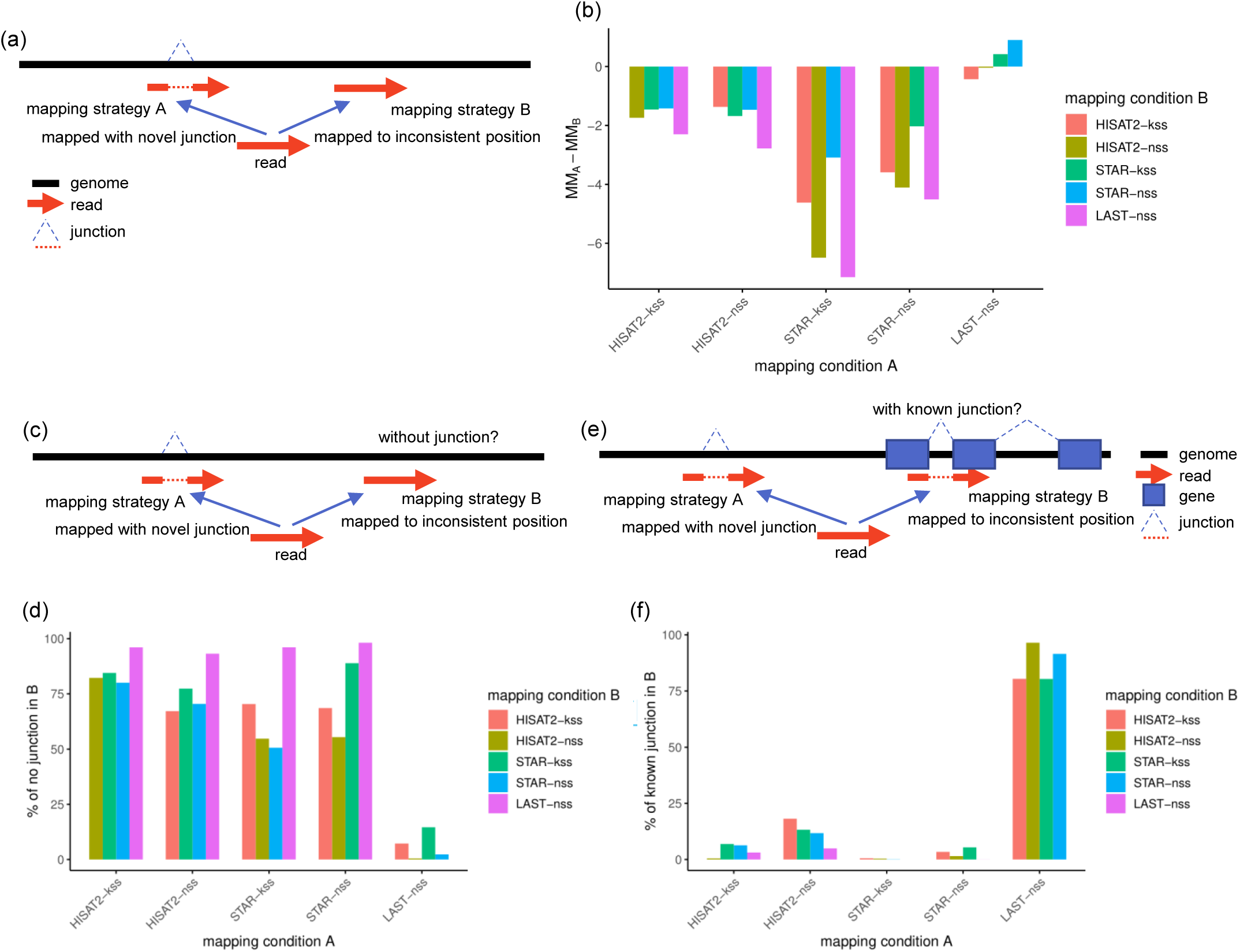
Inconsistently mapped reads with novel splice junctions. a) A model of an inconsistently mapped read with novel splice junctions. An inconsistently mapped read with novel splice junctions is one that contains novel splice junctions and that is mapped to 2 different locations by 2 different strategies (i.e., A and B). b) The difference in the mean number of mismatches between mapping strategies A and B. The *y*-axis indicates Mean(MM_Ai_ – MM_Bi_), where i is an inconsistently mapped read with novel splice junctions, MM_Ai_ is the number of mismatches of read i in mapping strategy A, and MM_Bi_ is the number of mismatches of read i for mapping strategy B. c, d) Models of inconsistently mapped reads with novel splice junctions. The models are reads with novel splice junctions that are mapped to one location by strategy A that mapped to an inconsistent position in strategy B when known splice junctions were absent (c) or included (d) in the analysis. e, f) The percentage of reads with novel splice junctions that were mapped by strategy A relative to all inconsistently mapped reads with novel splice junctions that were mapped by strategy B when known splice junctions were absent (e) or included (f) in the analysis.

### Same reads mapped to different positions

We obtained several cases in which replicate reads in the same sample mapped to inconsistent positions in HISAT-kss and HISAT-nss (Table S2). In addition, all strategies except STAR-kss sometimes mapped the same reads from different samples to different positions (Table S3). As an example, in the STAR-nss results, one sequence was mapped to the position “chr17:7307590, CIGAR: 1S47M” in two samples (ID: DRR021875 and DRR021877) but was mapped to the different location of “chr17:7307236, CIGAR: 1S33M353N14M” and was indicated to include a splice junction in a different sample (ID: DRR021876) (Table S4).

## Discussion

Many of the mapping software programs and options available for RNA-seq data are potentially applicable to the genome mapping of CAGE sequence data given that each CAGE sequence read is the 5□-end of an RNA. In this study, we showed that, with any of the strategies we evaluated, the mapping rate of CAGE data was approximately 60% to 70%, which is equivalent to standard RNA-seq mapping. In particular, the statistics regarding mapping results (e.g., mapped reads) were quite similar among HISAT2, STAR, and LAST. However, we noted small differences in the quantification of promoter expressions due to inconsistent alignments among mapping strategies. Although the correlations among mapping strategies were very high, the expression levels of at most ∼1.5% of FANTOM5 promoters differed significantly between different mapping strategies even when the same sequences were used. This difference was minimized by using a reference transcriptome, especially in STAR, but exacerbated depending on the mapping software applied.

When looking at individual alignments, we found significant differences among mapping strategies in regard to alignments to pseudogenes. The majority of reads inconsistently mapped to pseudogenes were mapped to protein-coding genes in another mapping condition (Fig. 3c,d). This finding suggests that these inconsistencies might be due to processed pseudogenes, which are similar to their parent genes but lack introns [23]. Generally identifying alignments to parent genes with short first exons is difficult due to the difficulty in detecting spliced alignments [5, 23, 24]; this situation leads to the preferred selection of alignments to processed pseudogenes with additional mismatches even if they are not expressed. Using a reference transcriptome can help to align reads over splice sites in parent genes, consistent with our observation that the number of expressed promoters associated with pseudogenes was lower in the strategies that included the reference transcriptome (i.e., HISAT2-kss and STAR-kss) than those that did not (i.e., HISAT2-nss and STAR-nss). However, even when we can correctly identify alignments both to processed pseudogenes and their parent genes, it is difficult to judge to which a sequence tag should be aligned. Given that nucleotides in processed pseudogenes are modified gradually and then mutated [23], we assumed that the potentially correct alignments were those with fewer mismatches, and we found that pseudogenes generally had more mismatches than their parent genes. Thus, we infer that most inconsistently mapped reads to pseudogenes might be incorrect, and the use of a reference transcriptome might facilitate choosing correct alignments between processed pseudogenes and parent genes with short first exons. In addition, the mean number of mismatches in the alignments to pseudogenes was lower in STAR than HISAT2 and LAST, indicating that pseudogene expression is likely more accurate when determined by using STAR than by using HISAT2 and LAST. We have not investigated why STAR yields better mapping results regarding pseudogenes than other mapping software programs, but the reason might be STAR’s better scheme for scoring of alignments that contain gaps [4].

Next, we found that mapping strategies differed in the detection of novel and known splice junctions. Except for LAST, the mean number of mismatches in alignments including novel splice junctions was generally smaller than their inconsistent reads, implying that the alignments with novel splice junction are more likely, as previously reported [4, 5]. Whether LAST appropriately mapped reads with novel splice junctions is difficult to assess, because the number of mismatches in alignments with novel splice junctions was similar between LAST and the other strategies. As expected, the use of a reference transcriptome did not influence the detection of novel splice junctions but contributed greatly to the detection of known splice junctions. These results suggest that it is better to use a reference transcriptome, given that mapping without it would miss more alignments of genes with short first exons. However, reference transcriptomes are unavailable or immature in many species. In these cases, HISAT2 is most appropriate for identifying novel splice junction candidates and likely would be useful for aggressive exploration of novel splice junctions. Compared with the other mapping strategies we evaluated, LAST tended to align reads with more mismatches rather than introducing novel junctions.

Sometimes in STAR 2-pass mapping, used in STAR-nss, replicate sequences were mapped to inconsistent positions in different samples, in which one read was mapped with splice junctions and another without splice junctions (Tables S3 and S4). Because these alignments have different 5□-ends, they hamper precise identification of TSS positions and their expression levels during CAGE analysis. During alignment postprocessing, we must remember that such cases might occur.

Although we specifically evaluated mapping software for CAGE data analysis, most of our results can be applied to other 5□-end sequencing protocols and even RNA-seq. Reads obtained by other 5□-end sequencing protocols—for example, TSS-seq and RAMPAGE—are similar to those obtained by CAGE. Because we did not focus on any points specific to CAGE protocols themselves, most of our findings can be applied to 5□-end sequencing protocols in general. Moreover, the different characteristics of the various genome mapping programs in regard to alignments to pseudogenes and with novel splice junctions could be critical for the analysis of reads obtained through RNA-seq protocols. Expression values of genes might be overestimated due to reads from their parent genes and underestimated due to undetected splice junctions.

## Conclusions

Overall, all strategies evaluated showed acceptable mapping rates and promoter rates. However, because CAGE reads tend to be short and because short first exons can influence mapping results, including a reference transcriptome in the mapping strategy is helpful. In particular, using a reference transcriptome can improve the mapping of reads between processed pseudogenes and their parent gene regions. In conclusion, mapping software and its conditions should be selected according to the particular research purposes, in light of the unique characteristics of each mapping strategy. For example, HISAT2 is particularly effective for discovering novel splice junctions, LAST yields high promoter rates, and STAR produces balanced results.

## Methods

### Sample information

The CAGE data for a human non-islet cell insulinoma cell line (TC-YIK) [17] were retrieved from DDBJ Sequence Read Archive (http://www.ddbj.nig.ac.jp/dra/) under the accession number DRA002420. The CAGE data comprised 93 samples, consisting of biological triplicates of 31 experimental conditions.

### Software and reference datasets

We used three mapping programs—HISAT2 (version 2.0.1-beta), LAST (version 852), and STAR (version 2.5.2a)—with the primary assembly of chromosomes 1 through 22, X, Y, and M for the human reference genome version GRCh38 (hg38). We used GENCODE human release 23 [1] as a reference transcriptome for HISAT2 and STAR. LAST has no options to use reference transcriptomes. For post-processing of mapping results, we used SAMtools (version 1.2) [25] and UCSC Genome Browser (version 356; http://hgdownload.cse.ucsc.edu/admin/exe/) [26, 27] software.

### Mapping and data processing

Before mapping, rRNA reads were removed by using rRNAdust (http://fantom.gsc.riken.jp/5/suppl/rRNAdust/) with a rRNA sequence file downloaded from UCSC Table Browser [28]. For each mapping strategy, we analyzed 93 samples; Table S1 shows the program version and options used. Reads with more than 5 soft-clipped bases at the aligned 5□ ends were flagged as unmapped and excluded from the further analysis. Mapping results in BAM format were processed to generate CAGE tag start sites (CTSS) files in BED format by extracting the 5□-end positions of uniquely mapped reads (the mapping quality > 20) [16]. The CTSS files were then converted to bigWig files by using the BedGraphTobigWig program in the UCSC Genome Browser software [26, 27].

### Data analysis

We counted all uniquely mapped reads as ‘mapped reads.’ The number of ‘expressed promoters’ was the number of hg38 FANTOM5 CAGE peaks [22] with more than 10 uniquely mapped reads. The ‘promoter rate’ was the number of reads within FANTOM5 CAGE peak regions divided by the total number of uniquely mapped reads. FANTOM5 CAGE peak annotation was obtained from the FANTOM5 website (http://fantom.gsc.riken.jp/5/datafiles/reprocessed/hg38_v4/extra/CAGE_peaks_annotation/hg38_liftover+new_CAGE_peaks_phase1and2_annot.txt). Categorization of promoters as driving the expression of a protein-coding gene, lincRNA, or pseudogene was based on the associated GENCODE transcripts in the annotation file; the number of expressed protein-coding genes, lincRNAs, or pseudogenes was the number of genes with at least one associated expressed promoter. Mapped reads containing splice junctions were retrieved as alignments whose CIGAR [25] values contained ‘N.’ Promoter-level expression was calculated by using the bigWigAverageOverBed program in the UCSC Genome Browser software [26, 27] using the FANTOM5 CAGE peaks [16] as a reference (“in.bed” of the program). In the differentially expressed promoter analysis, the edgeR package [29] in R version 3.3.3 (Bioconductor version 3.4) were used for biological triplicate samples (DRR021875, DRR021876, and DRR021877), and the q-value cutoff was 0.01. The hierarchical clustering was performed with the “hclust” function in R version 3.3.3.

### Novel splice junctions, pseudogenes, and mismatches

We defined a ‘novel splice junction’ as one where at least one side of the exon–intron junction was not matched with any (known) splice junction site in the GENCODE transcript annotation. The pseudogene list was obtained as the GENCODE genes labeled as “pseudogene.” The number of mismatches in an alignment was calculated as the sum of the number of “NM” tags in BAM files and the number of soft-clipped (HISAT2 and STAR) or hard-clipped (LAST) bases.

## List of abbreviations

CAGE: Cap Analysis of Gene Expression
CTSS: CAGE tag start sites
FANTOM: Functional Annotation of the Mammalian Genome
lincRNA: Long intervening non-coding RNA
nAnT-iCAGE: No-amplification non-tagging CAGE
RAMPAGE: RNA Annotation and Mapping of Promoters for Analysis of Gene Expression
TSS: Transcriptional start site

## Declarations

### Ethics approval and consent to participate

Not applicable.

### Consent for publication

Not applicable.

### Availability of data and material

The mapping result is available in our web site, https://genomec.gsc.riken.jp/gerg/owncloud/index.php/s/l6yCFRGdvLnhKkp.

### Competing interests

None declared

### Funding

This study was supported by KOFUKIN from MEXT through the RIKEN Center for Integrative Medical Sciences, RIKEN Preventive Medicine and Diagnosis Innovation Program, and RIKEN Centre for Life Science, Division of Genomic Technologies.

### Authors’ contributions

T.K. and H.K. conceived the study, designed the analyses, and coordinated the study. S.N. analyzed the data. S.N., T.K., and H.K wrote the manuscript. All authors read and approved the manuscript.

## Acknowledgements

We thank the FANTOM6 WG4 members of the FANTOM consortium for contributing to discussions of this study.

## Additional files

**Table S1** List of software options applied

**Table S2** Number of replicate sequences in the same sample that mapped to different positions

**Table S3** Number of replicate sequences in different samples that mapped to different positions

**Table S4** Example of a sequence that STAR-nss mapped to different positions in different samples

**Figure S1** Inconsistent reads mapped to pseudogenes.

The graph shows the percentage of inconsistently mapped reads between mapping strategies A and B in all reads mapped to pseudogenes in strategy A for each combination of mapping strategies A and B.

**Figure S2** Example of an inconsistently mapped read that was mapped to a pseudogene in one mapping strategy but to its parent gene in a different strategy.

Genomic views on UCSC Genome Browser are shown with the mapping results of DRR021875 by LAST-nss and STAR-kss. Graphs of CTSS and alignments of the read “DRR021875.14906” are shown for both mapping strategies. a) Genomic view around the TSS of RPL13. STAR-kss mapped more reads to this region than LAST-nss. For example, STAR-kss aligned the read “DRR021875.14906” to this region with a known splice junction of RPL13; LAST-nss did not map this read to this region. b) Genomic view around the TSS of RPL13P12, a pseudogene of RPL13P12. LAST-nss mapped more reads to this region than did STAR-kss. For example, LAST-nss aligned the read “DRR021875.14906” to this region without splice junctions; STAR-kss did not map this read to this region.

## Notes

https://genomec.gsc.riken.jp/gerg/owncloud/index.php/s/l6yCFRGdvLnhKkp

