## Supplementary Tables & Figures for "Assessment of mapping strategies for determining the 5□-end of mRNAs and long-noncoding RNAs with short read sequences"

Table S1 List of software options applied

<table>
<tr><th>mapping strategy</th><th>version</th><th>options</th></tr>
<tr><td>HISAT2-kss</td><td>version 2.0.1-beta</td><td>hisat2 -p <core> -t -x <genome_directory> -U <input> -S <output>.sam</td></tr>
<tr><td>HISAT2-nss</td><td>version 2.0.1-beta</td><td>hisat2 -p <core> --novel-splicesite-outfile <output>.splicesites.txt -t -x <genome_directory> -U <input> -S /dev/null<br>hisat2 -p <core> --novel-splicesite-infile <output>.splicesites.txt -t -x <genome_directory> -U <input> -S <output>.sam</td></tr>
<tr><td>LAST-nss</td><td>852</td><td>last-train -P<core> <genome_directory>/humandb <input>.fa > <ouput>.par<br>parallel-fasta "lastal -p <output>.par -d90 -D10 humandb | last-split -g humandb" < <input>.fa > <output>.maf</td></tr>
<tr><td>STAR-kss</td><td>2.5.2a</td><td>STAR  --readFilesIn <input>.fastq --runThreadN <core> --genomeDir <genome_directory> --outSAMtype BAM SortedByCoordinate --outBAMsortingThreadN <core> --outBAMcompression 6 --outFileNamePrefix <output></td></tr>
<tr><td>STAR-nss</td><td>2.5.2a</td><td>STAR --twopassMode Basic --readFilesIn <input>.fastq --runThreadN <core> --genomeDir <genome_directory> --outSAMtype BAM SortedByCoordinate --outBAMsortingThreadN <core> --outBAMcompression 6 --outFileNamePrefix <output></td></tr>
</table>

<core>: the number of core
<genome_directory>: the index for the reference genome
<input>: input file name
<output>: output file name

Table S2 Number of replicate sequences in the same sample that mapped to different positions

| | **DRR021875** | **DRR021876** | **DRR021877** |
|---|---|---|---|
| HISAT2-kss | 55 | 57 | 57 |
| HISAT2-nss | 78 | 85 | 73 |
| STAR-kss | 0 | 0 | 0 |
| STAR-nss | 0 | 0 | 0 |
| LAST-nss | 0 | 0 | 0 |

Table S3 Number of replicate sequences in different samples that mapped to different positions

| | **DRR021875 - DRR021876** | **DRR021876 - DRR021877** | **DRR021877 - DRR021875** |
|---|---|---|---|
| HISAT2-kss | 119 | 126 | 123 |
| HISAT2-nss | 1080 | 1089 | 1048 |
| STAR-kss | 0 | 0 | 0 |
| STAR-nss | 733 | 751 | 741 |
| LAST-nss | 936 | 167 | 812 |

Table S4 Example of a sequence that STAR-nss mapped to different positions in different samples

| | **DRR021875** | **DRR021876** | **DRR021877** |
|---|---|---|---|
| sequence | GAGGGCTGCTTGGTTGGTCAGTGGGGAGTCGGCGCCTGCGTACTAAGA | | |
| chr | chr17 | chr17 | chr17 |
| coord | 7307590 | 7307236 | 7307590 |
| strand | + | + | + |
| CIGAR | 1S47M | 3M353N45M | 1S47M |
| NM | 0 | 0 | 0 |

Figure S1

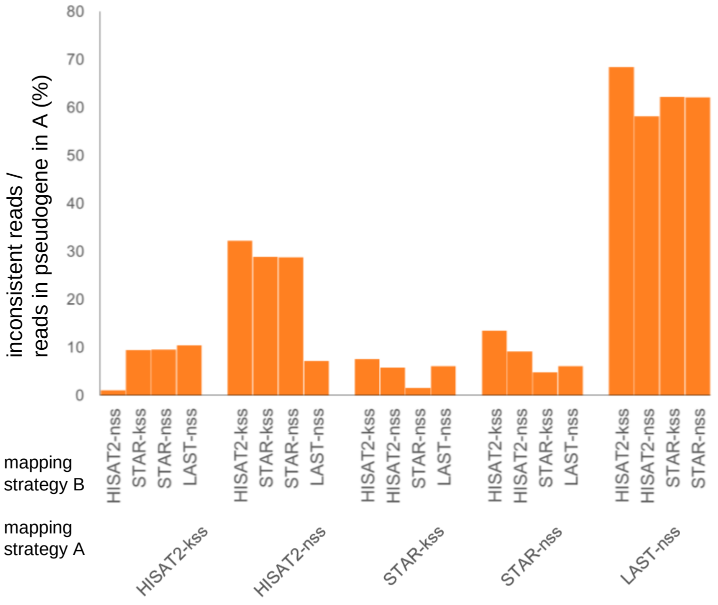
80
70
60
50
40
30
20
10
0
inconsistent reads /
reads in pseudogene in A (%)
HISAT2-nss
STAR-kss
STAR-nss
LAST-nss
HISAT2-kss
STAR-kss
STAR-nss
LAST-nss
HISAT2-kss
HISAT2-nss
STAR-nss
LAST-nss
HISAT2-kss
HISAT2-nss
STAR-kss
LAST-nss
HISAT2-kss
HISAT2-nss
STAR-kss
STAR-nss
mapping
strategy B
mapping
strategy A
HISAT2-kss
HISAT2-nss
STAR-kss
STAR-nss
LAST-nss

### Figure S2

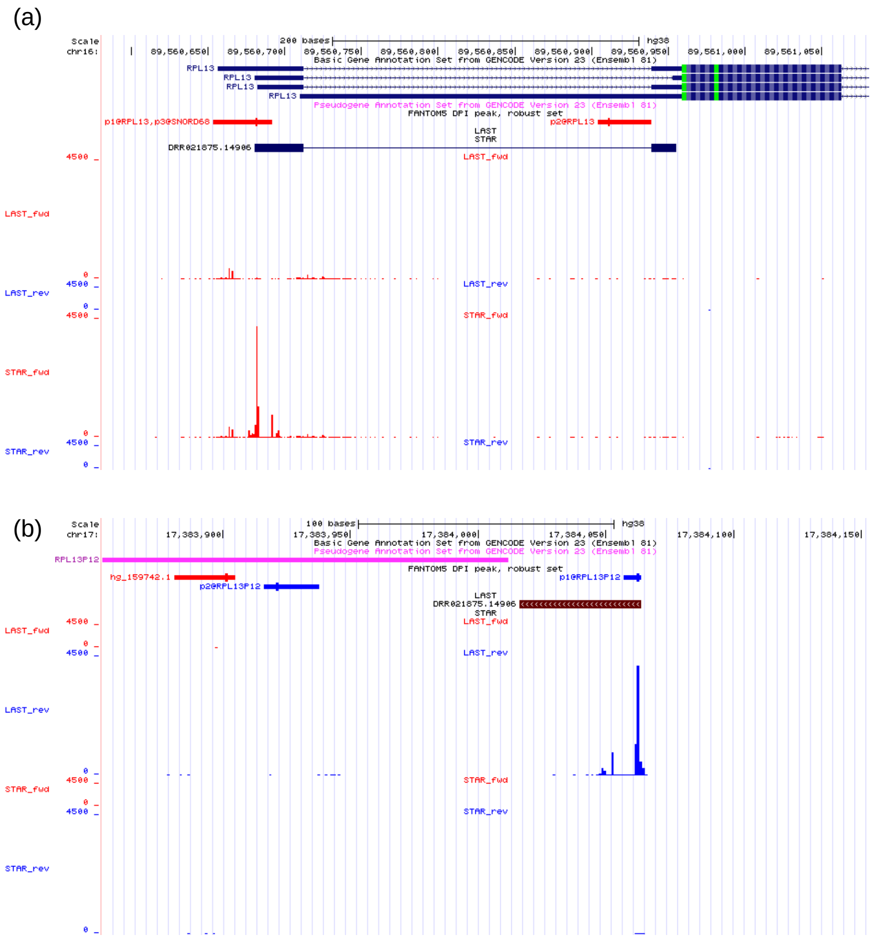
(a)
Scale
chr16:
200 bases
hg38
89,560,650
89,560,700
89,560,750
89,560,800
89,560,850
89,560,900
89,560,950
89,561,000
89,561,050
Basic Gene Annotation Set from GENCODE Version 23 (Ensembl 81)
RPL13
RPL13
RPL13
RPL13
Pseudogene Annotation Set from GENCODE Version 23 (Ensembl 81)
FANTOM5 DPI peak, robust set
p1@RPL13,p3@SNORD68
p2@RPL13
LAST
STAR
DRR021875.14906
LAST_fwd
LAST_rev
STAR_fwd
STAR_rev
4500
0
(b)
Scale
chr17:
100 bases
hg38
17,383,900
17,383,950
17,384,000
17,384,050
17,384,100
17,384,150
Basic Gene Annotation Set from GENCODE Version 23 (Ensembl 81)
Pseudogene Annotation Set from GENCODE Version 23 (Ensembl 81)
RPL13P12
FANTOM5 DPI peak, robust set
hg_159742.1
p2@RPL13P12
p1@RPL13P12
LAST
DRR021875.14906
STAR
LAST_fwd
LAST_rev
STAR_fwd
STAR_rev
4500
0
